# Capsule network for protein ubiquitination site prediction

**DOI:** 10.1101/2021.01.07.425697

**Authors:** Qiyi Huang, Jiulei Jiang, Yin Luo, Weimin Li, Ying Wang

**Author notes:** Corresporending author. (YL). These authors contributed equally to this work (Qiyi Huang, Jiulei Jiang). This author also contributed equally to this work. **Funding:** This project is supported by the National Key R&D Program of China (2018YFE0194000), National Nature Science Foundation of China (61762002), National Statistical Science Research Project (2020LY074). **Competing Interests:** The authors have declared that no competing interests exist.

## Abstract

Ubiquitination modification is one of the most important protein posttranslational modifications used in many biological processes. Traditional ubiquitination site determination methods are expensive and time-consuming, whereas calculation-based prediction methods can accurately and efficiently predict ubiquitination sites. This study used a convolutional neural network and a capsule network in deep learning to design a deep learning model, “Caps-Ubi,” for multispecies ubiquitination site prediction. Two encoding methods, one-of-K and the amino acid continuous type were used to characterize the sequence pattern of ubiquitination sites. The proposed Caps-Ubi predictor achieved an accuracy of 0.91, a sensitivity of 0.93, a specificity of 0.89, a measure-correlate-prediction of 0.83, and an area under receiver operating characteristic curve value of 0.96, which outperformed the other tested predictors.

## Introduction

Ubiquitination is an important posttranslational modification of proteins, consisting of the covalent binding of ubiquitin to a variety of cellular proteins. Ubiquitin was discovered in 1975 by Goldstein et al. [1]; it is a small protein composed of 76 amino acids [2]. Ubiquitination is the process of covalently binding the lysine of a substrate protein to the small ubiquitin molecule under the action of a series of enzymes. Three enzymes are involved in the process: E1 activation, E2 conjugation, and E3 ligation. Ubiquitination modification plays a very important role in basic reactions such as signal transduction, cell diseases, DNA repair, and transcription regulation [3–6]. Due to the important biological characteristics of ubiquitination, identifying potential ubiquitination sites helps to understand protein regulation and molecular mechanisms. Determining ubiquitination sites based on traditional biological experimental techniques such as mass spectrometry [7] and antibody recognition [8] is costly and time-consuming. Therefore, it is necessary to develop a calculation method that can accurately and efficiently recognize protein ubiquitination. In recent years, some calculation methods have been developed to predict potential ubiquitination sites. Huang et al. [9] used amino acid composition (AAC), a position weighting matrix, amino acid pair composition (AAPC), a position-specific scoring matrix (PSSM), and other information to develop a predictor called UbiSite using a support vector machine (SVM). Nguyen et al. [10] used an SVM to combine three kinds of information: AAC, evolution information, and AAPC to develop a predictor. Qiu et al. [11] developed a new predictor called “iUbiq-Lys” to apply to sequence evolution information and a gray system model. Chen et al. [12] also applied SVM to build a UbiProber predictor. Wang et al. [13] introduced physical–chemical attributes into an SVM to develop the ESA-UbiSite predictor. Radivojac et al. [14] developed the predictor UbPred using a random forest algorithm. Lee et al. [15] developed UbSite using efficient radial basis functions. All of those machine learning-based methods and predictors have promoted the development of ubiquitination site prediction research and achieved good prediction performance. However, most of them rely on artificial feature selection, which may lead to imperfect features [16], and their datasets are small despite the large volume of accumulated biomedical data.

Deep learning, the most advanced machine learning technology, can handle large-scale data well. It has multilayer networks and nonlinear mapping operations, which can fit the complex structure of data well. In recent years, deep learning has been developed rapidly [16] and has been successfully applied in various fields of bioinformatics [17,18]. Some methods based on deep learning have been used for ubiquitination site identification. For example, Fu et al. [19] applied one-hot and composition of k-spaced amino acid pairs encoding methods to develop DeepUbi with text-CNN. Liu et al. [20] used deep transfer learning methods to develop the DeepTL-Ubi predictor for multispecies ubiquitination site prediction. He et al. [21] established a multimodel predictor using one-hot, physical–chemical properties of amino acids, and a PSSM.

Although various ubiquitination site predictors and tools have been developed, there are still some limitations, and their accuracy and other performance elements must be further improved. In this paper, a deep learning model, “Caps-Ubi,” is proposed that uses a capsule network for protein ubiquitination site prediction. In Caps-Ubi, the protein fragments are first passed through one-of-K and amino acid continuous methods to encode them. Then three convolutional layers and the capsule network layer are used as a feature extractor to obtain the functional domains in the protein fragments and finally to get the prediction result. Relative to existing tools, the prediction performance of Caps-Ubi is a significant improvement. Researchers could use the predictor to select potential ubiquitination candidate sites and do experiments to verify them, which will reduce the range of protein candidates and save time.

## Materials and methods

### Benchmark dataset

The ubiquitination dataset came from the largest online protein lysine modification database, PLMD 3.0, which contains 20 protein lysine modifications. The database has 53,501 proteins and 284,780 protein lysine modification sites, including 25,103 proteins and 121,742 ubiquitination sites. To eliminate errors caused by homologous sequences, we used CD-HIT [22] to filter out homologous sequences with sequence similarities greater than 40%. We obtained 12,100 proteins and 54,586 ubiquitination sites, which were used as a positive sample set. Based on those annotated sequences, 427,305 nonubiquitinated sites were extracted from the proteins as a negative sample set, and CD-HIT-2D [23] was used to filter out homologous sequences within the positive sample set that were greater than 50%. To establish a balanced training model, we randomly selected the same data as the positive sample set and selected 90% of it as the training and validation sets and 10% as the independent test set. Finally, 53,999 data on ubiquitination sites and 50,315 data on nonubiquitination sites were obtained. The final data division is shown in Table 1.

**Table 1.** Data of protein ubiquitination sites

| Dataset | No. of positive data | No. of negative data |
| --- | --- | --- |
| Training | 44,214 | 44,214 |
| Validation | 4,913 | 4,913 |
| Testing | 5,459 | 5,459 |

### Input sequence coding

The coding method directly determines the quality of its prediction results; a good feature can extract the correlation between the ubiquitination feature and the targets from peptide sequences [24]. After encoding the protein sequence, the sequence information is converted into digital information, and then deep learning is done on it. In this study, two methods were used to encode the amino acid sequence around the protein ubiquitination site; namely, one-of-K encoding and amino acid continuous encoding.

#### One-of-K encoding

The one-of-K encoding method was adopted for protein fragments, and each protein fragment was encoded into an *m* × *k* 2D matrix, where *m* is the number of amino acids in each sequence— that is, the length of the input sequence—and *k* is the type of amino acid. There are 20 kinds of common amino acids. When the length of the input sequence did not reach the window length, it was filled in with a “-” on the left or right side of the protein fragment and was treated as another amino acid, so each sequence consisted of 21 amino acids.

#### Continuous coding of amino acids

The continuous amino acid coding method [25] was proposed by Venkatarajan; the coding uses 237 physical-chemical properties to quantitatively characterize 20 amino acids. They used five main components to characterize the changes in 237 physica-chemical properties of amino acids. In this paper, each amino acid is represented by a 6D vector, wherein the first 5D represents the five principal components as shown in Table 1 of [25], the last 1D represents the gap in the input protein fragment with a length of *m*. The gap is represented by a dash”-”, meaning that when the sequence length does not reach the window length, the bit is coded as 1; otherwise, it is 0. Finally, each protein fragment is coded into an *m* × 6 2D matrix. This continuous coding scheme can comprehensively consider the physical and chemical properties of protein amino acids and has a smaller dimension than that of one-of-K coding. The smaller input dimension will lead to a relatively simple network structure, which is beneficial to avoid overfitting.

### Capsule network

In a CNN, the pooling layer can extract valuable information from the data, but some location information is lost [26]. Also, a CNN outputs scalar values in neurons, and the information represented by scalar neurons is limited and cannot reflect the spatial position relation of the internal features of the neural network. To solve the problems of scalar neurons, in 2017 Hinton proposed a deep learning architecture called a capsule network [27]. The main building module of a capsule network is the capsule [28], which is a set of neuron vectors. The length of the capsule represents the probability of the existence of an entity; the longer the capsule, the greater the probability,and the direction of the capsule represents the state of the entity. The capsule network provides a unique and powerful deep learning building block that can better model the complex relations within a neural network. A CNN uses scalar input activation functions, such as the rectified linear activation function ReLU, a sigmoid, and a tanh, and the capsule network uses an activation function called a squash. The calculation equation is

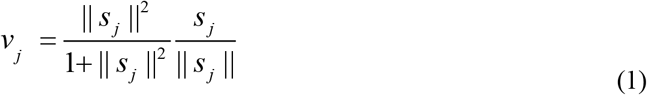

where *v* _*j*_ is the output of capsule *j*, and *s* _*j*_ is the weighted sum of the input vectors of capsule *j*. This function compresses the vector length to the interval [0,1], which can be regarded as a kind of compression and reallocation of the vector length. In addition to the first-layer capsule network, the input of the capsule *s* _*j*_ is obtained by the weighted sum of the prediction vector (*û* _*j* |*i*_) located in the lower-layer capsule, and the prediction vector (*û* _*j* |*i*_) is passed through the lower layer. The capsule is calculated by multiplying its output (*u* _*i*_) and the weight matrix (*w* _*ij*_):

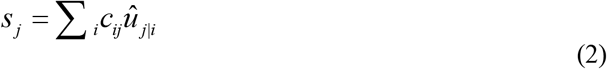

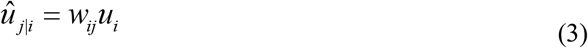

where *c*_*ij*_ is the coupling coefficient, which is obtained by a softmax transformation from *b*_*ij*_; its calculation equation is

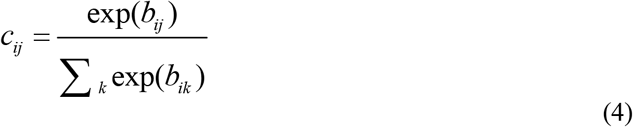

In Eq. (4), the sum of the coupling coefficients of all capsules and capsule *i* in the previous layer is 1. The coupling coefficient is obtained through a dynamic routing mechanism; the pseudocode is as follows:

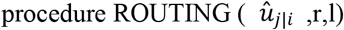

for all capsules *i* in layer l and capsules *j* in layer (l + 1): *b*_*ij*_ 0.

for *r* iterations do:

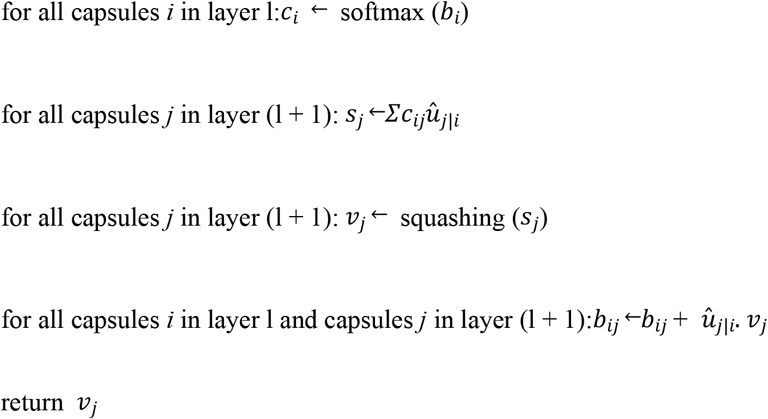

The loss function of the capsule network is the margin loss function, and the calculation equation is

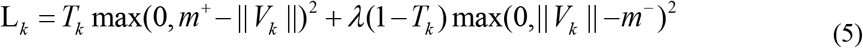

where *K* is the number of categories, *T* _*K*_ is the real label ubiquitinated to 1 and nonubiquitinated to 0, ||*V* _*k*_ || is the output length of the *k*th capsule, which is the probability of predicting the *k*th class. The boundary *m* ^+^ is 0.9, which is a penalty for false positives, and the lower boundary *m* ^―^ is 0.1, which is a penalty for false negatives. *λ* is a proportional coefficient of 0.5, which is used to control the loss caused when some categories do not appear, to prevent the capsule vector length of all categories from being reduced in the early stage of training,and the total loss is the sum of the losses of *K* categories.

### Architecture design

As shown in Figure 1, the structure of the proposed model contains two identical subnetworks that process one-of-21 and amino acid continuous encoding modes. After training in their respective network model, the two models merge the features as the final output. Each subnetwork consists of the same three 1D convolutional layers (Conv1, Conv2, Conv3) and a capsule network layer. The first convolutional layer (Conv1) of the network is a 1D convolution kernel, which comprises 256 convolution kernels with a size of 1 and a step size of 1 that use the ReLU activation function. A convolution kernel with a length of 1 first appears in the Network in Network [29]; a convolution kernel with a length of 1 can reduce the complexity of the model and can make the network deeper and wider. Applied in this study, it acted as a feature filter and could pool features in two encoding modes. The second convolutional layer, Conv2, is a conventional convolutional layer with 256 1D convolution kernels with a length of 7 and a step size of 1, which functions as a local feature detector to extract the protein sequence input and convert it to corresponding local features. Conv2 is understood as the functional domain characteristics of the protein, and its output is used as the input of the next layer, Conv3. The third convolutional layer, Conv3, has 256 1D convolution kernels with a size of 11 and a step size of 1. The activation function used is ReLU and a dropout mechanism with a random deletion rate of 0.3. The dropout mechanism is used to prevent the model from overfitting and to increase the generalization ability of the model. These two convolutional layers are used to increase the feature representation ability of the capsule network and convert the original features of protein fragments into more advanced and abstract features. Then the local features of Conv2 are used as the input of the PrimaryCapsule network layer. The dimension of each capsule in the PrimaryCapsule is 8, the step size is 1, the convolution kernel length is 20, and the squash activation function is used. The last layer of LabelCapsule is a capsule with a dimension of 10, which is used to represent the two states of the input protein fragment: the input sequence is ubiquitination site or non-ubiquitination site, and finally the output of the two subnetworks are merged as the final prediction result.

**Figure 1.**
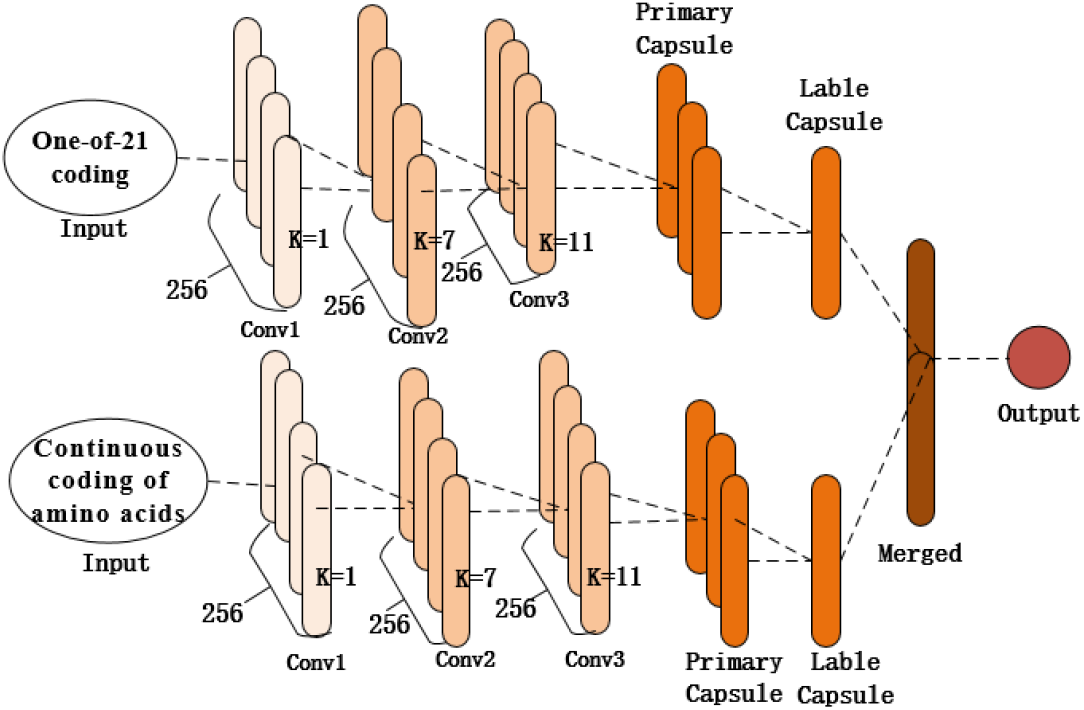
Network structure structure of the proposed model

### Model training

For model training, we used the Adam[30] optimization algorithm. Adam can automatically adjust the learning rate of the parameters, improve the training speed, and improve the stability of the model. The learning rate was 0.003, the first-order estimated exponential decay rate was 0.9, and the exponential decay rate estimated by the second moment was 0.999. The dynamic routing mechanism was consistent with that in the original paper [26]. The number of routing iterations was 3, and the boundary loss function was used as the loss function of the model. The boundary loss function form is shown in Eq. (5). and the number of model training iterations was 50 epochs. The deep learning framework used by this model was Keras 2.1.4. Keras is a highly modular deep learning framework based on Theano and written in Python; it supports both CPU and GPU. The programming language was Python 3.5, and the model was trained and tested on a Windows 10 system equipped with an Nvidia RTX 2060 GPU.

## Result

### Model evaluation and performance indicators

A confusion matrix is a visual display tool used to evaluate the quality of classification models. Each row of the matrix represents the actual condition of the sample, and each column represents the sample condition predicted by the model. There are four values in the matrix, as shown in the following equations, where *FN* is the number of false negatives, *FP* is the number of false positives, *TN* is the number of true negatives, and *TP* is the number of true positives. The following indicators based on the confusion matrix are usually used to evaluate the prediction of the model performance:

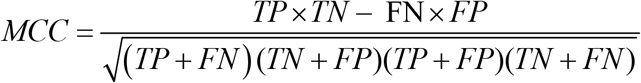

Among them, *S*_*n*_ stands for sensitivity, which is the evaluation of the prediction performance of negative samples; *S*_*p*_ is the specificity, which is the evaluation of the prediction performance of positive samples; *Acc* is the accuracy, which is the evaluation of the accuracy of the model; and *MCC* is the Matthew’s correlation coefficient, which is the overall evaluation of the model. The receiver operating characteristic (ROC) curve and the area under the curve (AUC) for the ROC curve are usually used to evaluate the pros and cons of binary classifiers: the larger the AUC value, the better the model performance.

### Experimental results

First, we did many experiments on the selection of the window size of protein fragments. Because the correlation information between amino acids had a direct effect on the prediction results, we needed to determine an appropriate window size. Previous studies directly used empirical values such as 21, 33, or 49. However, different data models and classifiers tend to have different window sizes [31]. Therefore, a window length of *n* was selected from a range of 21 to 75, and we did a series of experiments with the different window lengths. For each window length, we encoded all training data into two input modes and trained their respective subnetworks. According to the prediction results of the validation set, we selected each appropriate window size. Figure 2 shows the performance of various window sizes in one-of-21 and amino acid continuous encoding modes.

**Figure 2.**
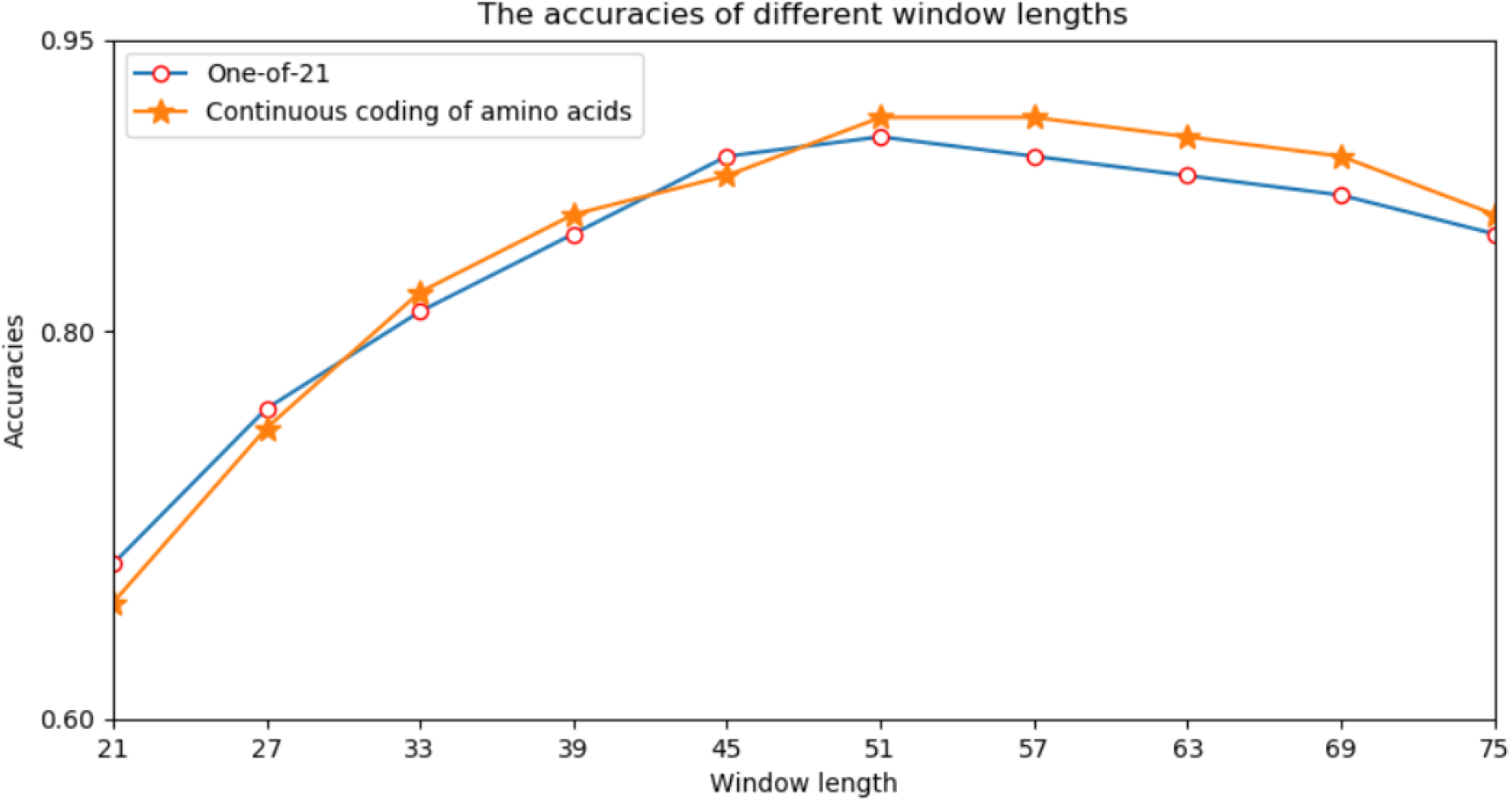
Accuracy of the verification set for various window lengths

In Figure 2, the abscissa represents the window length, and the ordinate represents the accuracy of the model. It can be seen from Figure 3 that when the window length was 51, the two encoding modes had the highest accuracy. Therefore, we set the window length of this model to 51.

**Figure 3.**
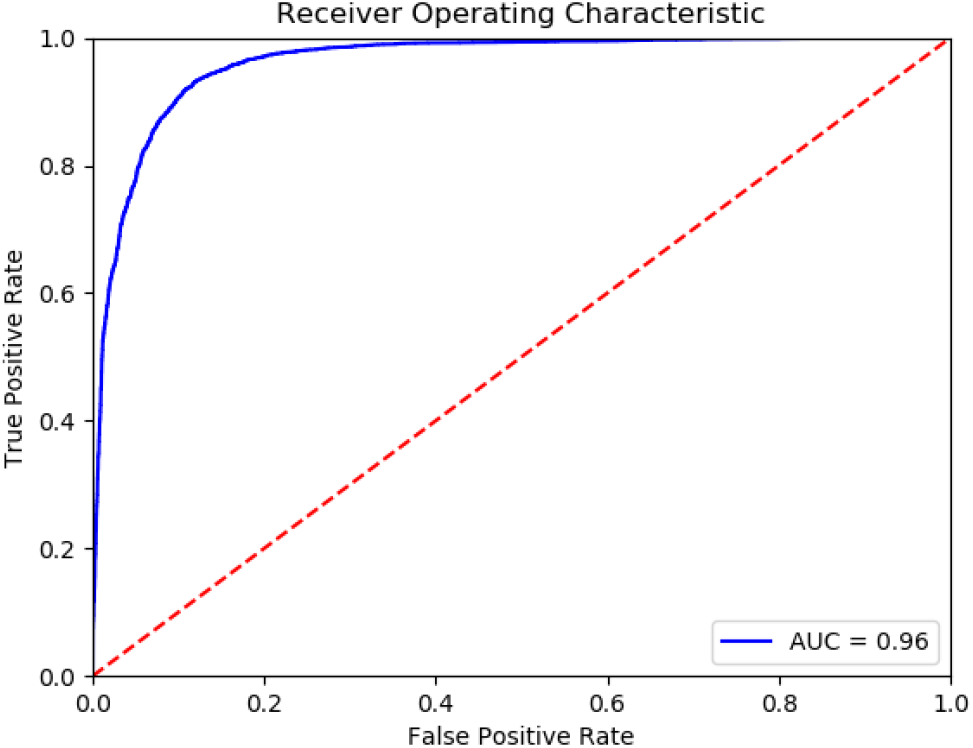
Receiver operating characteristic curve of Caps-Ubi on the test set

To compare the performance of the model under different encoding schemes, we compared the capsule network and the CNN with similar hierarchical structures of capsule networks and the same training set size. The CNN structure replaced only the PrimaryCapsule layer with the Conv3 layer. We set the LabelCapsule layer to a 128 × 1 fully connected layer. The comparison results are shown in Table 2.

**Table 2.** Comparison of various coding schemes

| Feature | Model | Acc (%)<sup>1</sup> | Sn (%)<sup>2</sup> | Sp (%)<sup>3</sup> | AUC<sup>4</sup> | MCC<sup>5</sup> |
| --- | --- | --- | --- | --- | --- | --- |
| One-of-21 | CapsNet | 89.51 | 93.70 | 85.31 | 0.96 | 0.80 |
|  | CNN | 84.93 | 86.39 | 82.93 | 0.93 | 0.70 |
| Amino acid continuous | CapsNet | 90.06 | 91.88 | 88.23 | 0.96 | 0.80 |
|  | CNN | 83.83 | 85.25 | 82.41 | 0.91 | 0.68 |
| One-of-21 and amino acid continuous | CapsNet | 90.47 | 93.66 | 87.27 | 0.96 | 0.81 |
|  | CNN | 84.67 | 82.62 | 86.72 | 0.93 | 0.70 |

<sup>1</sup>Accuracy of the model<sup>2</sup>Sensitivity of the model<sup>3</sup>Specificity of the model<sup>4</sup>Area under curve<sup>5</sup>Matthew's correlation coefficient

From Table 2, it can be concluded that the capsule network’s accuracies were 5.39%, 7.43%, and 6.85% percentage points higher than those of CNN under the one-of-21, amino acid continuous, and combined one-of-21 and amino acid continuous types, indicating that the capsule network internally expressing the hierarchical relation modeling aspect has more advantages than CNN. Among them, the performance under the combined one-of-21 and amino acid continuous encoding modes is the best on the capsule network: this proposed Caps-Ubi model achieved an accuracy, sensitivity, specificity, area under curve, and Matthew’s correlation coefficient of 91.23%, 93.11%, 89.34%, 0.96, 0.83 respectively. The proposed Caps-Ubi was obtained from balanced data. The ROC curve of Caps-Ubi on the test set is shown in Figure 3, which shows that it was very close to the real situation.

When we used balanced data to train the model on an experimentally verified ubiquitination dataset and a nonubiquitination dataset [19], the ratio of positive peptides and negative peptides was 1:8, so we tested Caps-Ubi using natural-distribution data. The test results are shown in Table 3. According to the test results, the performance was slightly worse than that under the balanced data.

**Table 3.** Results of testing Caps-Ubi under natural-distribution data

| Protein fragment | Acc (%)<sup>1</sup> | Sn (%)<sup>2</sup> | Sp (%)<sup>3</sup> | AUC<sup>4</sup> | MCC<sup>5</sup> | Positive-negative ratio |
| --- | --- | --- | --- | --- | --- | --- |
| 1,000 | 53.75 | 0.08 | 0.99 | 0.70 | 0.19 | 1:8 |
| 10,000 | 53.30 | 0.12 | 0.95 | 0.59 | 0.12 | 1:8 |

<sup>1</sup>Accuracy of the model<sup>2</sup>Sensitivity of the model<sup>3</sup>Specificity of the model<sup>4</sup>Area under curve<sup>5</sup>Matthew's correlation coefficient

### Comparison with other methods

In the past 10 years, many researchers have contributed to the prediction and research of protein ubiquitination sites. We compared the proposed model with other sequence-based prediction tools. The corresponding data and results are shown in Table 4, which shows that the performance of the Caps-Ubi model exceeded that of the best-performing deep learning model DeepUbi and several other prediction models. The accuracy, sensitivity, specificity, area under curve, and Matthew’s correlation coefficient of Caps-Ubi were 2.36, 3.31, 1.24, 0.05, and 0.05 respectively percentage points higher than those of DeepUbi.

**Table 4.**
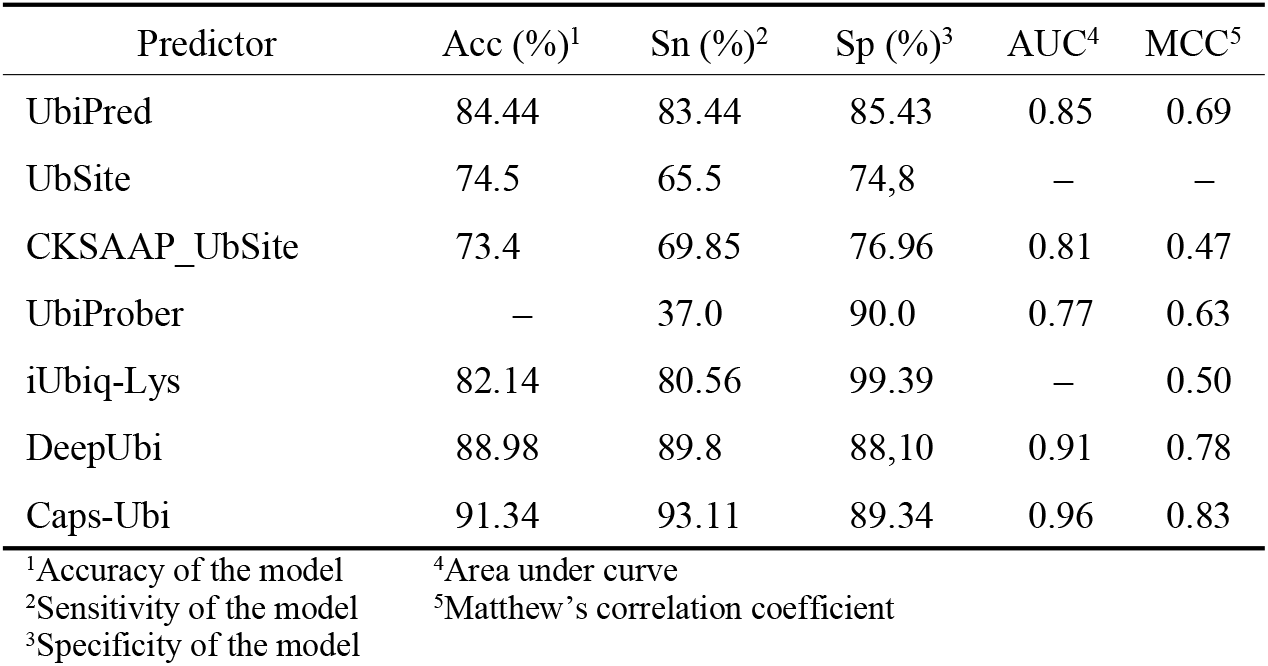
Proposed Caps-Ubi compared with other methods

## Conclusion and outlook

In this paper, a new deep learning model for predicting protein ubiquitination sites is proposed, using one-of-K and amino acid continuous coding modes. We used the largest available protein ubiquitination site dataset, and the experimental results above verify the effectiveness of this model. The operation of the model has four main steps: encoding protein sequences, constructing convolutional layers, constructing a capsule network layer, and constructing an output layer. The capsule network introduces a new building block for deep learning. Relative to CNN, the capsule network, which uses a dynamic routing mechanism to update parameters, requires more training time, but the time required for prediction is similar. The capsule network can also characterize the complex relations among amino acids in various sequence positions and can explore the internal data distribution related to biochemical significance. The proposed Caps-Ubi prediction tool will facilitate the sequence analysis of ubiquitination and can also be used to identify other posttranslational modification sites in proteins. In the future, we will study other features that may better extract sample attributes to construct deeper models.

## References

1. Goldstein G, Scheid M, Hammerling U, Schlesinger DH, Niall HD, Boyse EA. Isolation of a polypeptide that has lymphocyte-differentiating properties and is probably represented universally in living cells. Proc Natl Acad Sci U S A. 1975;72:11–15.

2. Wilkinson KD. The discovery of ubiquitin-dependent proteolysis. Proc Natl Acad Sci U S A. 2005;102:15280–15282.

3. Hicke L, Schubert HL, Hill CP. Ubiquitin-binding domains. Nat Rev Mol Cell Biol. 2005;6:610–621.

4. Hicke L. Protein regulation by monoubiquitin. Nat Rev Mol Cell Biol. 2001;2:195–201.

5. Pickart CM. Ubiquitin enters the new millennium. Mol Cell. 2001;8:499–504.

6. Haglund K, Dikic I. Ubiquitylation and cell signaling. EMBO J. 2005;24:3353–3359.

7. Peng J, Schwartz D, Elias JE, et al. A proteomics approach to understanding protein ubiquitination. Nat Biotechnol. 2003;21:921–926.

8. Gentry MS, Worby CA, Dixon JE. Insights into Lafora disease: malin is an E3 ubiquitin ligase that ubiquitinates and promotes the degradation of laforin. Proc Natl Acad Sci U S A. 2005;102(24):8501–8506.

9. Huang CH, Su MG, Kao HJ, Jhong JH, Weng SL, Lee TY. UbiSite: incorporating two-layered machine learning method with substrate motifs to predict ubiquitin-conjugation site on lysines. BMC Syst Biol. 2016;10 Suppl 1(Suppl 1):6.

10. Nguyen VN, Huang KY, Huang CH, Lai KR, Lee TY. A New Scheme to Characterize and Identify Protein Ubiquitination Sites. IEEE/ACM Trans Comput Biol Bioinform. 2017;14:393–403.

11. Qiu WR, Xiao X, Lin WZ, Chou KC. iUbiq-Lys: prediction of lysine ubiquitination sites in proteins by extracting sequence evolution information via a gray system model. J Biomol Struct Dyn. 2015;33:1731–1742.

12. Chen X, Qiu JD, Shi SP, Suo SB, Huang SY, Liang RP. Incorporating key position and amino acid residue features to identify general and species-specific Ubiquitin conjugation sites. Bioinformatics. 2013;29:1614–1622.

13. Wang JR, Huang WL, Tsai MJ, Hsu KT, Huang HL, Ho SY. ESA-UbiSite: accurate prediction of human ubiquitination sites by identifying a set of effective negatives. Bioinformatics.2017;33:661–668.

14. Radivojac P, Vacic V, Haynes C, et al. Identification, analysis, and prediction of protein ubiquitination sites. Proteins. 2010;78(2):365–380.

15. Lee TY, Chen SA, Hung HY, Ou YY. Incorporating distant sequence features and radial basis function networks to identify ubiquitin conjugation sites. PLoS One. 2011;6:e17331.

16. Wang D, Zeng S, Xu C, et al. MusiteDeep: a deep-learning framework for general and kinase specific phosphorylation site prediction. Bioinformatics. 2017;33:3909–3916.

17. Shaw D, Chen H, Jiang T. DeepIsoFun: a deep domain adaptation approach to predict isoform functions. Bioinformatics. 2019;35(15):2535–2544.

18. Sun, D., Wang, M., Feng, H., & Li, A.. (2018). Prognosis prediction of human breast cancer by integrating deep neural network and support vector machine: Supervised feature extraction and classification for breast cancer prognosis prediction. 2017 10th International Congress on Image and Signal Processing, BioMedical Engineering and Informatics (CISP-BMEI). IEEE.

19. Fu H, Yang Y, Wang X, Wang H, Xu Y. DeepUbi: a deep learning framework for prediction of ubiquitination sites in proteins. BMC Bioinformatics. 2019;20:86.

20. Liu Y, Li A, Zhao XM, Wang M. DeepTL-Ubi: A novel deep transfer learning method for effectively predicting ubiquitination sites of multiple species. Methods. 2020;S1046-2023(20)30156-0.

21. He F, Wang R, Li J, Bao L, Xu D, Zhao X. Large-scale prediction of protein ubiquitination sites using a multimodal deep architecture. BMC Syst Biol. 2018;12(Suppl 6):109.

22. Huang Y, Niu B, Gao Y, Fu L, Li W. CD-HIT Suite: a web server for clustering and comparing biological sequences. Bioinformatics. 2010;26:680–682.

23. Huang CH, Su MG, Kao HJ, Jhong JH, Weng SL, Lee TY. UbiSite: incorporating two-layered machine learning method with substrate motifs to predict ubiquitin-conjugation site on lysines. BMC Syst Biol. 2016;10 Suppl 1(Suppl 1):6.

24. Plewczynski D, Tkacz A, Wyrwicz LS, Rychlewski L. AutoMotif server: prediction of single residue post-translational modifications in proteins. Bioinformatics. 2005;21:2525–2527.

25. Venkatarajan M S, Braun W. New quantitative descriptors of amino acids based on multidimensional scaling of a large number of physical–chemical properties[J]. Molecular modeling annual, 2001, 7(12):445–453.

26. Dombetzki LA. An overview over capsule networks. Network Architectures and Services 2018.

27. Sabour S, Frosst N, Hinton G E. Dynamic Routing Between Capsules[J]. 2017.

28. Hinton,G.E. et al. (2011) Transforming Auto-encoders. International Conference on Artifificial Neural Networks. Springer, Finland, pp. 44–51.

29. Lin M., Chen Q., Yan S. Network in network[J]. arXiv preprint 1312.4400, 2013:

30. Kingma, D. and Ba, J. (2014) Adam: a method for stochastic optimization, arXiv preprint 1412.6980

